# Motility-Induced Phase Separation Mediated by Bacterial Quorum Sensing

**DOI:** 10.1101/2023.04.01.535124

**Authors:** Wesley J. M. Ridgway, Mohit P. Dalwadi, Philip Pearce, S. Jonathan Chapman

## Abstract

We study motility-induced phase separation (MIPS) in living active matter, in which cells interact through chemical signalling, or quorum sensing. In contrast to previous theories of MIPS, our multiscale continuum model accounts explicitly for genetic regulation of signal production and motility. Through analysis and simulations, we derive a new criterion for the onset of MIPS that depends on features of the genetic network. Furthermore, we identify and characterise a new type of oscillatory instability that occurs when gene regulation inside cells promotes motility in higher signal concentrations.

## INTRODUCTION

Chemical signals control cell motility to regulate self-organised patterning in living systems, from tissue morphogenesis and wound healing to cancer [1]. In bacterial populations, chemical signalling, or quorum sensing (QS), drives self-organisation by promoting population-level behaviours such as biofilm formation and swarming motility [2, 3]. Bacterial QS systems have been engineered to connect directly to genes controlling motility in synthetic genetic networks, enabling the generation of tunable patterns *in vitro* [4–6]. However, we lack fundamental understanding of how cell-level features, such as gene-regulatory networks, determine emergent population-level pattern formation in living active matter.

Minimal physical models of active and living matter typically consist of physically interacting self-propelled particles. Despite their simplicity, such models can display complex emergent dynamics, such as motility-induced phase separation (MIPS) [7, 8]. This phenomenon is caused by a self-trapping mechanism whereby particles experience reduced motility at high densities owing to their interactions, eventually leading to dense macroscopic clusters of immotile particles coexisting with a dilute phase of motile particles [7]. Similar phase transitions have been observed in active matter systems with agents that interact through flows [9–13], morphogens [14], social interactions [15, 16], chemotaxis [17–21], and electrostatic torques [22]. In the past, models for chemically interacting particles have been placed within this framework by representing signals indirectly through a density-dependent motility [7, 23–25] or an effective physical force [26, 27], but it is not known how far these approaches accurately represent chemical signalling in living matter.

More recently, minimal models of active matter have been supplemented with a concentration field of signalling molecules that mediates the orientation [21, 28–34] or motility [5, 35–37] of cells or particles. Such models have demonstrated that, broadly, repression of motility by intercellular signals tends to promote variations in cell density [5, 36, 38–41], in line with models of physically interacting particles. However, these existing models often represent gene regulation and chemical signalling using coarse-grained or effective terms, rather than accounting for gene-regulatory kinetics explicitly. Therefore, it is not known in general how the properties of gene-regulatory networks connect to the characteristics of emergent patterns in living active matter.

Here, we develop a multiscale continuum model of chemically interacting particles or cells. Our theory systematically accounts for intracellular processes through careful treatment of the population’s gene regulation and phenotype (i.e. motility). Thereby, we connect population-level patterns with the gene-regulatory network inside cells and intercellular QS signalling. We derive a criterion for MIPS mediated by QS in terms of the properties of the gene-regulatory network and discover a new route to MIPS via genetic regulation of tumbling frequency. We clarify that it is only consistent to approximate chemically mediated interactions by a density–dependent motility in the limit of fast chemical timescales; in general, one must account for chemical timescales to predict the onset of QS-induced MIPS. In general, the density dependence is nonlinear, in contrast to physically interacting active Brownian particles (ABPs) [42–44]. Finally, we identify and explain a new type of oscillatory instability that occurs when cell motility is promoted, rather than repressed, in higher concentrations of QS signal. This new instability does not occur in active matter systems in which the interactions are purely physical.

## MODEL SETUP

We start by imposing rules at the cell level, before systematically deriving a continuum model from the cell dynamics. We consider a population of *N* chemically interacting bacteria in two dimensions, and neglect physical interactions to focus on the role of chemical signalling.

Each cell is denoted by the index *i* and is located at position **x**_*i*_ with an internal chemical concentration *u*_*i*_(*t*), which represents a gene-regulatory protein or transcription factor. The internal chemical obeys the kinetics

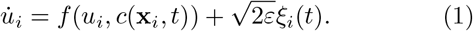

Here, *f* characterises the gene-regulatory network (GRN), and depends on the internal chemical concentration *u*_*i*_ and the local concentration field *c* of autoinducer (AI), which permeates the population. Although our upscaling is general, we later define *f* for definiteness. The last term in (1) represents the stochastic behaviour inherent to chemical reactions; *ξ*_*i*_ represents zero-mean Gaussian white noise and *ε* its magnitude. We work in the limit of small magnitude noise in the gene-regulatory kinetics. We include this small noise term for two reasons. Firstly, biochemical reactions are intrinsically stochastic, especially if the reagents are present in small amounts [45]. Secondly, this term regularizes the continuum equations and, as we show below, the system is singular in the limit *ε* → 0.

In terms of cell motility, we assume that the cells undergo both active motion and passive diffusion. The active component is characterised by active Brownian motion [46, 47] and run-and-tumble dynamics. Between tumbles, the particle positions **x**_*i*_ therefore obey

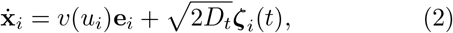

where *v*(*u*_*i*_) is the gene-regulated self-propulsion speed and *D*_*t*_ a passive translational diffusion coefficient associated with the zero-mean Gaussian white noise **ζ**_*i*_. The cell orientations are described by the vector **e**_*i*_ = (cos *ϕ*_*i*_, sin *ϕ*_*i*_)^*T*^ where the angle *ϕ*_*i*_ undergoes diffusion with no directional bias due to the active Brownian component:

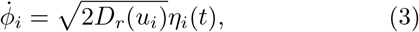

where *D*_*r*_ is a rotational diffusion coefficient associated with the zero-mean Gaussian white noise *η*_*i*_. The active motility parameters *D*_*r*_, *v*, and the tumble rate *γ* are gene-regulated, which is modelled by allowing a dependence on *u*_*i*_. Such regulation may arise naturally [3, 48] or synthetically [4–6].

We derive continuum equations from the individual-based equations of motion (1)–(3) for a large population of cells using standard methods (see Section A of the Supplemental Material [49], c.f. [44, 50–52]). In the continuum equations, the intracellular chemical concentration *u* becomes an independent variable. That is, the population-level equations are structured in terms of the GRN inside cells; we refer to these multiscale equations as GRN-structured.

The continuum GRN-structured cell density *n*(**x**, *u, t*) satisfies

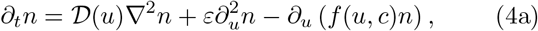

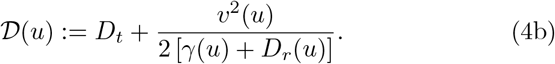

Here, 𝒟 (*u*) is the effective gene-regulated diffusion coefficient. The second term on the RHS of (4a) accounts for the stochastic component of the kinetics. The third term on the RHS of (4a) codifies the GRN through an advection of the structured cell density in the *u*-coordinate. The second term on the RHS of (4b) is the contribution arising from the run-and-tumble and active Brownian dynamics. The physical cell density *ρ*(**x**, *t*) is related to the GRN-structured cell density *n* through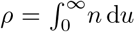. In deriving equations (4), we assume that the gene-regulatory kinetics and cell diffusion occur on timescales much longer than the tumbling and reorientation timescales, in line with biological parameter estimates [36, 53], and consider lengthscales much larger than the cell persistence length (see Section A of the Supplemental Material [49]).

Positive feedback is a canonical component of quorum-sensing GRNs, present in many bacterial species [2, 53]. We pose the following specific functional form for *f* which incorporates positive feedback:

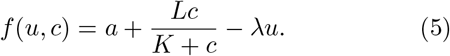

Here, *a* represents a constant base production rate of the intracellular chemical *u*, and *λ* is a natural decay rate. The second term on the RHS of (5) represents the production of *u* induced by the local AI concentration *c*, which constitutes one half of the positive feedback loop illustrated in Figure 1a. This term saturates at a maximal rate of *L* and has a ‘threshold’ activation at *c* = *K* where the production rate is half-maximal.

**FIG. 1:**
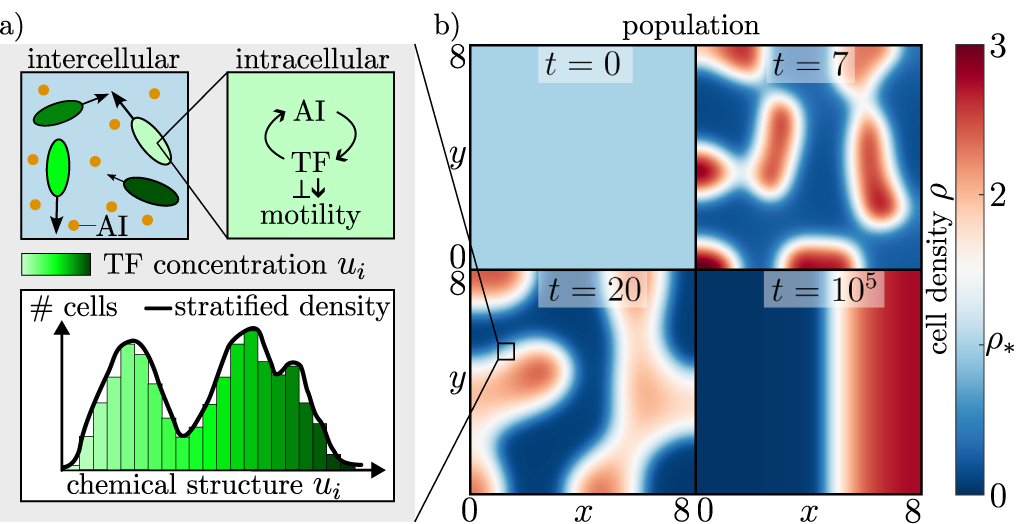
a) Schematic illustration of our multiscale model of chemically interacting cells. We derive a continuum population model (right), that retains the genetic structure of the population (grey box, left) through chemical stratification. The motility of individual cells depends on their internal chemical concentration. The transcription factor (TF) can either repress or promote motility. b) Snapshots in time of the cell density from an initially homogeneous state (AI concentration profile is similar). See also Video S1 in the Supplemental Material [49].

The other half of the positive feedback loop in the QS circuit involves the AI concentration field *c*(**x**, *t*). We assume that AI diffuses passively with coefficient *D*_*c*_, decays with rate *β*, and is generated through cell secretion at a rate *α*(*u*). Thus, we have

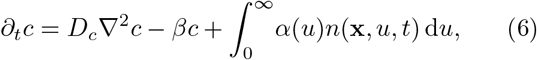

where the final term on the RHS is the continuum secretion term. This term is non-local in *u* as it encodes the contribution from all internal concentrations in a locally averaged region of space. We emphasize that positive feedback is only present in the QS circuit when the secretion rate *α*(*u*) is non-constant across internal concentrations *u*. For simplicity, we pose a secretion rate that is proportional to the internal concentration, i.e.

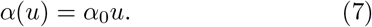

Finally, we assume that the cell population is confined to a rectangular domain Ω, imposing no flux boundary conditions on the boundaries *∂*Ω. Hence,

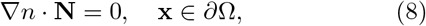

where **N** is the unit normal on *∂*Ω. To ensure physical (i.e. non-negative, bounded) concentrations, we also impose no flux at *u* = 0 and as *u* →*∞*. These conditions correspond to

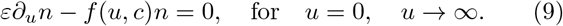

Our continuum model therefore consists of the governing equations (4)–(7), and the boundary conditions (8)– (9). The multiscale nature of our model is illustrated schematically in Figure 1a. We verify the predictions of our theory in the following section via numerical simulations of the governing equations, using the open-source finite-element library oomph-lib [54] (see Section E of the Supplemental Material [49]).

## INSTABILITY OF THE UNIFORM EQUILIBRIUM STATE

To investigate the emergence of MIPS in our model, we search for instabilities in the *spatially* uniform equilibrium state, emphasizing that this is not *chemically* uniform in general. To this end, we perform a linear stability analysis of the governing equations (4)–(9) to derive an instability criterion for the onset of MIPS.

We derive the steady uniform solution by directly integrating (4a) and imposing the boundary conditions (9). The spatially uniform equilibrium is then given by

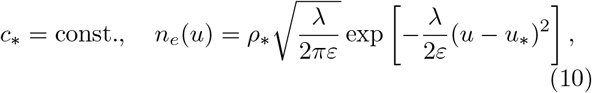

where the steady AI concentration *c*_*\**_ and mean internal concentration *u*_*\**_ are defined through the unique positive solution of the algebraic system

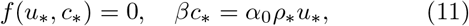

where 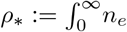 d*u* represents the uniform cell density.

The analysis is singular in the limit *ε* → 0, since the equilibrium density *n*_*e*_ (10) formally tends to a Dirac delta function centred at *u* = *u*_*\**_, representing identical internal concentrations in each cell. To analyse this singular perturbation problem, we perform a WKBJ-like asymptotic approximation and explicitly factor out the singular exponential term. This reduces the analysis to a regular perturbation problem (see Section B of the Supplemental Material [49]).

We perform the linear stability analysis by substituting small perturbations of the form

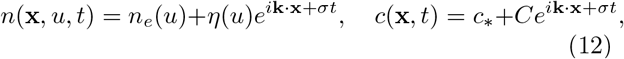

into the governing equations (4)-(6) and linearizing the result. Here, **k** denotes the wavenumber and *σ* the growth rate of the small perturbations. This yields the dispersion relation (see Section B of the Supplemental Material [49])

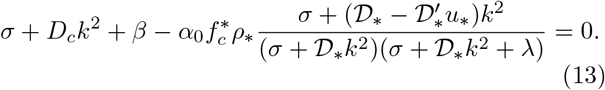

Here *k* := |**k**| while 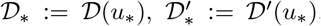, and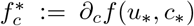 . The spatially uniform steady-state (10) is unstable if Eq. (13) has any root σ with Re(σ) *>* 0. We refer to 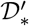 as the motility response since it characterise how the motility responds to changes in internal concentration. We find that there are two types of instability depending on the sign of 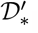.

### Case I: Internal chemical represses motility

The first type of instability occurs when higher internal concentrations reduce motility 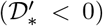, analogous to classic MIPS where higher cell densities reduce motility through physical interaction.

We derive the following instability criterion by substituting *σ* = 0 into (13) and rearranging:

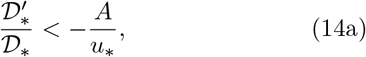

where we define

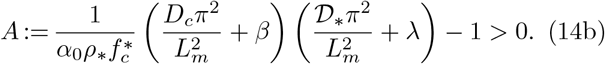

Here, *L*_*m*_ is the largest side length of the domain, and we use *k* = *π/L*_*m*_ since the longest wavelength mode is always the first to lose stability as 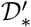 is decreased from zero (see Section B of the Supplemental Material [49]). However, the mode with the largest growth rate, i.e. largest value of Re(*σ*), typically corresponds to an intermediate wavelength, as shown in Fig 2.

**FIG. 2:**
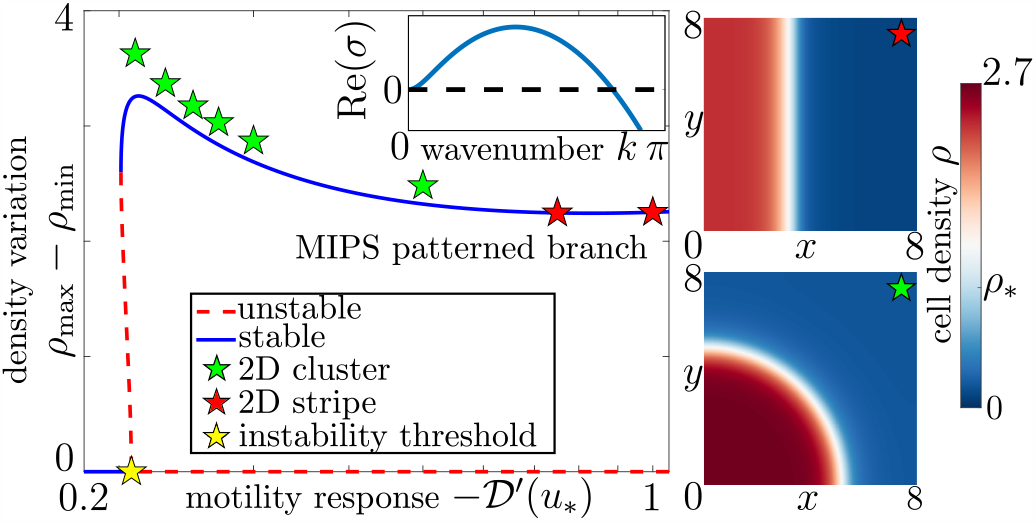
Bifurcation diagram in 1D with 2D steady-state stripe (cluster) patterns shown as red (green) stars, respectively (see right panels and Video S2 in the Supplementary Material). The trivial branch corresponds to (10) and the bifurcating branch is the 1D MIPS-patterned state (computed from steady-state versions of (4)–(7)). The bifurcation point (yellow star) is indistinguishable from that predicted from (14). Inset: dispersion relation from linear theory. Parameter values are given in Table S1 of the Supplemental Material [49].

Eq. (14) is the first key result of our paper. Qualitatively, it states that a spatially uniform population of chemically interacting cells begins to form clusters when the cellular motility (characterized by the diffusion co-efficient 𝒟_*\**_) is sufficiently repressed in response to perturbations in the internal concentrations. The constant *A*, defined in (14b), encodes the required strength of repression in terms of the chemical timescales and genetic structure of the population. The mechanism driving the instability is shown schematically in Figure 4a.

A key feature of the instability characterised by (14) is that any of the gene-regulated active motility parameters in (4b) can trigger MIPS, not just the self-propulsion speed *v*(*u*). This is in contrast to physical MIPS for which the analog of (14) is given by the criterion [7]:

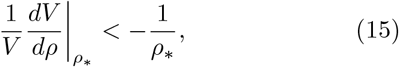

where *V* (*ρ*) is the effective density-dependent propulsion speed in physical active matter. A similar phenomenon occurs for active (velocity) fluctuations where rotational diffusion affects the onset of MIPS, but is not the underlying cause [50]. The difference between (14) and (15) is that (14) accounts for the timescales of chemical diffusion and gene-regulated motility parameters. As such, we expect that the classical result (15) should be recovered when both the AI concentration and gene-regulatory kinetics equilibriate quickly. We clarify in Sections B and D of the Supplemental Material [49] that in order to recover (15), the chemical timescales need to be fast not just by comparison to the cell diffusion timescale but also by comparison to the tumbling and reorientation timescale, which necessitates modifying the upscaling procedure used to obtain (4). In this very-fast-chemical-timescale limit, non-constant rotational diffusion or tumbling frequency cannot cause MIPS, consistent with (15).

The dynamics of the cell density *ρ* are illustrated in Figure 1b, clearly exhibiting MIPS. In Figure 2 we compute steady-state branches that bifurcate from the uniform state, and show that the bifurcation point predicted by our analysis in (14) agrees well with numerically computed branches. The unstable spatial modes from the dispersion relation *σ*(*k*), determined from (13), are illustrated in Figure 2. The spatial mode with the largest growth rate measures the cluster sizes that initially form from the uniform state.

### Case II: Internal chemical promotes motility

Surprisingly, our linear stability analysis predicts a new type of instability when higher internal concentrations promote motility 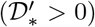. Since physical inter-particle forces act to reduce motility as density increases, this new instability would not be induced in populations with purely physical interactions. Chemically interacting populations do not have this restriction; GRNs connected to intercellular signalling can increase cell motility directly in e.g. bacterial swarming where QS regulates flagella assembly [48], and indirectly by e.g. controlling biosurfactant production [3].

Mathematically, this instability occurs when *σ* is purely imaginary, i.e. via a Hopf bifurcation. The analogue of (14a) for this instability is given by (see Section C of the Supplemental Material [49]):

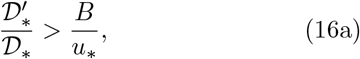

where we define

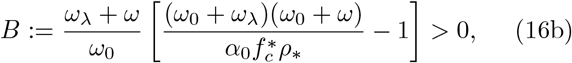

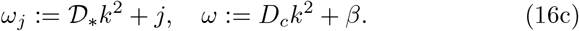

Eq. (16) is the second key result of our paper. It quantifies the condition under which a uniform population of chemically interacting cells undergo spatio-temporal density oscillations. Qualitatively, the population becomes unstable when the cellular motility is sufficiently promoted in response to perturbing the internal concentrations. The constant *B*, defined in (16b), encodes information about the chemical timescales and genetic structure of the population. Similar to the criterion (14), the instability can be triggered if any of the active motility parameters in (4b) are non-constant. In contrast to (14), the longest wavelength mode, in general, is not the first to become unstable, as can be seen by comparing the dispersion relations for the two instabilities in Figures 2 and 3c. We show the type of spatio-temporal dynamics arising from this instability in Figure 3a. The oscillations tend to a limit cycle resembling a standing wave pattern, which bifurcates from the uniform state as illustrated by Figure 3b.

**FIG. 3:**
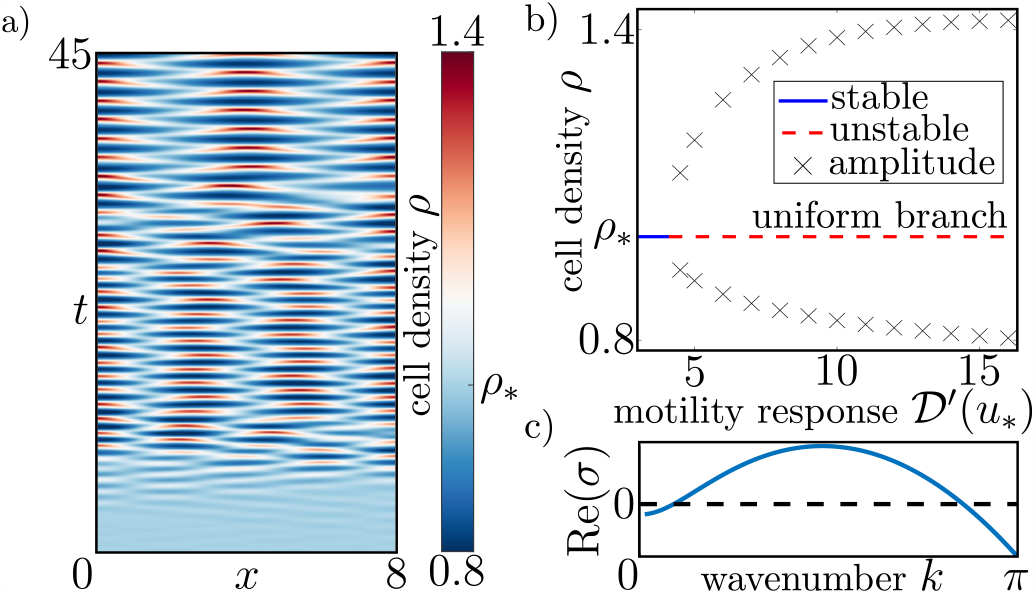
a) Oscillatory patterning in 1D showing the formation of periodic spatio-temporal oscillations resembling standing waves (see also Video S3 in the Supplemental Material). b) Bifurcation diagram showing the amplitude of density oscillations. c) Dispersion relation from the linear theory.

The mechanism driving this new oscillatory instability is an effective time delay between local density fluctuations and changes in motility, illustrated schematically in Figure 4b. Oscillatory patterns are known to arise in chemotactic active matter where effective time-delays between changes in density and orientation [21] or chemoattractant [29] drive the instability. Here, the time delay between density and motility is caused by finite timescales in the QS circuit. To understand the mechanism physically, consider a region of locally higher cell density. As the density fluctuation begins to relax, the AI concentration field increases locally in response to the higher density, which leads to locally higher internal concentrations due to the reaction kinetics. Owing to the chemically regulated motility (*u*), the cells with higher internal concentrations experience a higher motility. As the local density returns to the equilibrium value, the locally higher motility persists and the region begins to deplete of cells, thereby forming a region of lower density. This forms the first half of the periodic cycle, with the second half having equivalent reasoning. The oscillations decay if the time delay is too short, which can be seen mathematically by considering the behaviour of *B* as the delay tends to zero, i.e. in the limit of fast chemical timescales (in which *a*∼ *L* ∼*λ*∼ *α*_0_ ∼*β* are large). From (16b), it can be shown that *B*→*∞* in this limit and hence the criterion (16a) cannot be satisfied, favouring a stable uniform colony.

**FIG. 4:**
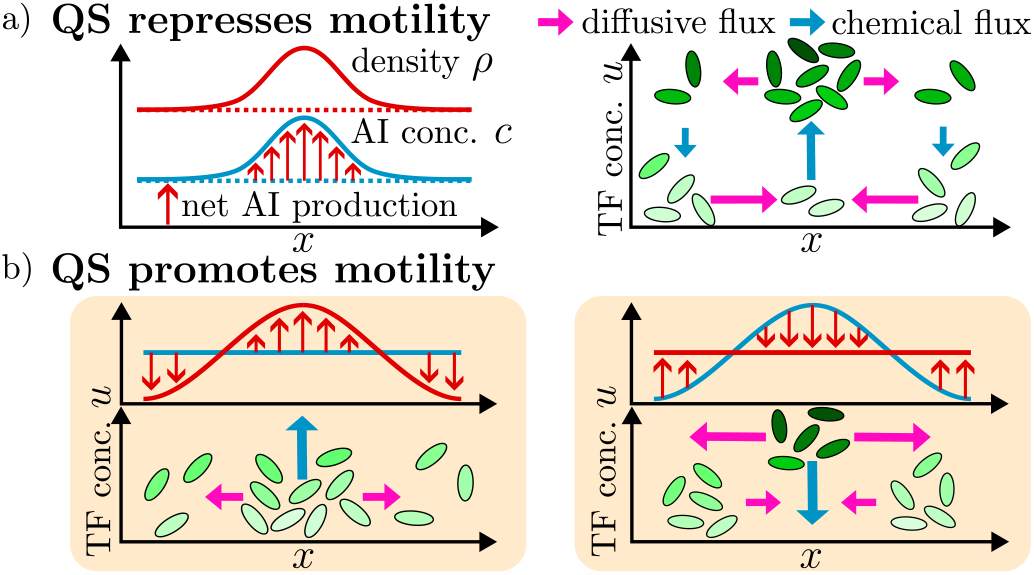
Schematic illustrations of the instability mechanisms. a) Small density/AI perturbation from the unstable uniform equilibrium in Case I. b) Periodic spatio-temporal oscillations in Case II.

Our results demonstrate that explicit modelling of gene regulation in cell populations is key to understanding MIPS in living active matter. Our GRN–structured model allows for a direct link between cell-level genetic processes and macroscale pattern formation – in principle, the model can be simplified via an internal mean-field formulation of the governing equations (4)–(7), but this is only quantitatively accurate near the spatially uniform equilibrium (Supplemental Material [49]). More fundamentally, our theory predicts that gene-regulated tumbling frequency alone can cause MIPS, in contrast to classical physical MIPS. Additionally, gene regulation that promotes motility in higher signal concentrations is required for the oscillatory instability (Hopf bifurcation) that we identify here; this instability is absent in systems with purely physical interactions between particles. Our continuum model is appropriate for large cell populations, which are common in many natural [3] and synthetic [5] biological systems. In future work it would be interesting to explore the predictions of agent-based simulations of the microscopic model (2)–(3), (5)–(6), especially for small cell populations.

## ACKNOWLEDGEMENTS

The authors thank Devi Prasad Panigrahi and Andrew Hazel for helpful discussions. Furthermore, the authors would like to acknowledge the use of the University of Oxford Advanced Research Computing (ARC) facility in carrying out this work, http://dx.doi.org/10.5281/zenodo.22558, as well as the UCL Mathematics Department for use of the high performance cluster. W. J. M. R. is grateful for financial support in part from an NSERC PGSD scholarship and in part from Somerville College. M. P. D. is supported by the UK Engineering and Physical Sciences Research Council [EP/W032317/1]. P. P. is supported by a UKRI Future Leaders Fellowship [MR/V022385/1].

