## Supplemental Material for "Motility-Induced Phase Separation Mediated by Bacterial Quorum Sensing"

October 4, 2023

#### A Derivation of the Governing Equation for the Chemically Structured Cell Density

In this section we derive the governing equation (4) from the equations of motion (1)–(3). The derivation uses the approach of [1,2] in considering a continuum formulation of the individual-based dynamics. This involves using the many-body Fokker-Planck equation to formulate the equations of motion in terms of a probability density. We then derive a governing equation for the one-particle probability density from which we obtain (4) by considering an appropriate macroscopic scaling regime for the length and timescales.

We start with an equivalent description of (1)–(3) in terms of the joint probability density $\psi(\{\mathbf{x}_i, \phi_i, u_i\}, t)$ of the population being in the state $\{\mathbf{x}_i, \phi_i, u_i\}_{i=1}^N$ at time $t > 0$. The many-body Fokker-Planck equation describes the evolution of $\psi$; it reads

$$\partial_t \psi = \sum_{i=1}^N \nabla_i \cdot \left(-v(u_i)\mathbf{e}_i + D_t \nabla_i\right) \psi + \sum_{i=1}^N D_r(u_i) \partial^2_{\phi_i} \psi + \sum_{i=1}^N \left( \frac{\gamma(u_i)}{2\pi} \int_0^{2\pi} \psi \, \mathrm{d}\phi_i - \gamma(u_i)\psi \right)$$
$$+ \sum_{i=1}^N \partial_{u_i} \left[\varepsilon \partial_{u_i} \psi - f(u_i, c(\mathbf{x}_i, t))\psi\right], \quad \mathbf{x}_i \in \Omega, \quad u_i \in \mathbb{R}_+, \quad \phi_i \in [0, 2\pi). \tag{A.1}$$

The respective fluxes in the first set of brackets are due to self-propulsion at speed $v$ in the direction $\mathbf{e}_i = (\cos\phi_i, \sin\phi_i)^T$, and translational diffusion with associated diffusion coefficient $D_t$. The rotational diffusion component of the active Brownian motion has a diffusion coefficient of $D_r$ and is captured by the terms in the second sum. The respective terms in the third sum describe tumbles into, and out of orientation angle $\phi_i$, respectively. The non-local term implicitly assumes (for simplicity) that tumbling events are instantaneous and that the new orientation angle is chosen from a uniform distribution over $[0, 2\pi)$. The time intervals between tumbles are assumed to be exponentially distributed with rate parameter $\gamma$. Finally, the fluxes in the last sum encode the kinetics $f$ of the GRN with a stochastic component of strength $\varepsilon$. The gene-regulatory kinetics depend on the local concentration of AI $c(\mathbf{x}, t)$. We assume that the population is isolated, imposing no flux boundary conditions in $x_i$ and $u_i$, as well as periodic boundary conditions in $\phi_i$.

The next step in obtaining a continuum formulation of (1)–(3) is to derive an effective equation that governs the evolution of the one-particle probability density $\psi_1$. Since the particles are identical, we WLOG consider the conditional probability for the first particle, i.e.

$$\psi_1(\mathbf{x}, \phi, u, t) := N \int \psi \, \mathrm{d}\phi_2 \ldots \mathrm{d}\phi_N \, \mathrm{d}\mathbf{x}_2 \ldots \mathrm{d}\mathbf{x}_N \, \mathrm{d}u_2 \ldots \mathrm{d}u_N, \tag{A.2}$$

where the subscript '1' has been dropped on the independent variables. The equation for $\psi_1$ is obtained by integrating (A.1) over the positions, orientations, and internal concentrations of particles $2, \ldots, N$. The result is

$$\partial_t \psi_1 = \nabla \cdot (D_t \nabla \psi_1 - v\mathbf{e}\psi_1) + D_r \partial_\phi^2 \psi_1 + \frac{\gamma}{2\pi}\int_0^{2\pi} \psi_1 \,\mathrm{d}\phi - \gamma\psi_1 + \partial_u \left[\varepsilon \partial_u \psi_1 - f(u, c(\mathbf{x}, t))\psi_1\right], \tag{A.3}$$

where we have used the no flux boundary conditions in $\mathbf{x}_i$ and $u_i$ as well as the periodic boundary conditions in $\phi_i$.

Next, we derive governing equations for the chemically structured cell density $n$ and orientation field $\mathbf{p}$, defined as the zeroth and first angular moments of $\psi_1$. In particular,

$$n(\mathbf{x}, u, t) := N \int \psi_1 \,\mathrm{d}\phi_1, \tag{A.4a}$$

$$\mathbf{p}(\mathbf{x}, u, t) := N \int \mathbf{e}\psi_1 \,\mathrm{d}\phi_1, \tag{A.4b}$$

where $\mathbf{e} = (\cos\phi, \sin\phi)^T$. We consider large populations of cells ($N \gg 1$) for which a continuum macroscale description is appropriate for the moments $n$ and $\mathbf{p}$. This allows the secretion term in the governing equation (6) in the main text to be written in terms of $n$. The governing equation for $n$ is found by integrating equation (A.3) over the orientation while the governing equation for $\mathbf{p}$ is found by first multiplying by $\mathbf{e}$ and then integrating. The equation for $n$ reads

$$\partial_t n = -\nabla \cdot \mathbf{J}_p - \partial_u J_c, \qquad \mathbf{J}_p = -D_t \nabla n + v(u)\mathbf{p}, \qquad J_c = -\varepsilon \partial_u n + f(u, c)n, \tag{A.5a}$$

where $\mathbf{J}_p$ is the 'physical' component of the flux due to translational diffusion and active motion. The 'chemical' flux $J_c$ encodes the GRN (c.f. the last sum in (A.1)). The orientation field satisfies

$$\partial_t \mathbf{p} = D_t \nabla^2 \mathbf{p} - \frac{v(u)}{2}\nabla n - \omega \mathbf{p} + \varepsilon \partial_u^2 \mathbf{p} - \partial_u \left(f(u, c)\mathbf{p}\right) + \mathbf{M}[\psi_1], \qquad \omega(u) = D_r(u) + \gamma(u). \tag{A.5b}$$

The second and third terms on the RHS of (A.5b) state that local orientational order is induced by density gradients and naturally dissipates at the reorientation and tumbling rate $\omega$. Here $\mathbf{M}[\psi]$ is a higher-order angular moment of $\psi_1$ given by

$$\mathbf{M}[\psi_1] = -v(u) \int_0^{2\pi} \left(\mathbf{e}\mathbf{e}^T - \frac{\mathbf{I}}{2}\right) \nabla \psi_1 \,\mathrm{d}\phi, \tag{A.6}$$

where $\mathbf{I}$ is the identity matrix.

Before introducing a physiologically relevant scaling regime, we non-dimensionalise (A.5). To this end, we introduce the dimensionless quantities

$$\tilde{\mathbf{x}} = \frac{\mathbf{x}}{\mathcal{L}}, \quad \tilde{t} = \frac{t}{\mathcal{T}}, \quad \tilde{u} = \frac{u}{\mathcal{U}}, \quad \tilde{n}(\tilde{\mathbf{x}}, \tilde{u}, \tilde{t}) = \frac{n(u, \mathbf{x}, t)}{\mathcal{N}}, \quad \tilde{\mathbf{p}}(\tilde{\mathbf{x}}, \tilde{u}, \tilde{t}) = \frac{\mathbf{p}(\mathbf{x}, u, t)}{\mathcal{P}}, \quad \tilde{c} = \frac{c}{\mathcal{C}}, \tag{A.7a}$$

as well as the dimensionless parameters

$$\tilde{\omega}_a(\tilde{u}) = \frac{\omega(u)}{\Omega}, \quad \tilde{v}(\tilde{u}) = \frac{v(u)}{V} \quad \tilde{D}_t = \frac{D_t \mathcal{T}}{\mathcal{L}^2}, \quad \tilde{D}_c = \frac{D_c \mathcal{T}}{\mathcal{L}^2}, \quad \tilde{\alpha_0} = \frac{\alpha_0 \mathcal{T}\mathcal{U}^2\mathcal{N}}{\mathcal{C}}, \quad \tilde{\beta} = \mathcal{T}\beta, \tag{A.7b}$$

where tildes denote dimensionless quantities. Here $\mathcal{L}$ and $\mathcal{T}$ are respective characteristic length and timescales, while $\mathcal{U}$ and $\mathcal{C}$ are characteristic concentrations of internal chemical and AI. The quantities $\mathcal{N}$ and $\mathcal{P}$ are typical values of the structured cell density and orientation field magnitude, respectively. We have also introduced the quantities $\Omega$ and $V$, which are typical values of the cell tumbling/reorientation rate $\omega$ and velocity $v$. We additionally define the dimensionless parameters

$$\chi := \Omega\mathcal{T}, \quad \Gamma := \lambda\mathcal{T}, \tag{A.7c}$$

and non-dimensionalise the kinetic parameters in (1) and (5) as

$$\tilde{\varepsilon} = \frac{\varepsilon}{\lambda \mathcal{U}^2}, \quad \tilde{a} = \frac{a}{\lambda \mathcal{U}}, \quad \tilde{L} = \frac{L}{\lambda \mathcal{U}}, \quad \tilde{K} = \frac{K}{\mathcal{C}}, \quad \tilde{f}(\tilde{u}, \tilde{c}) = \tilde{a} + \frac{\tilde{L}\tilde{c}}{\tilde{K} + \tilde{c}} - \tilde{u}. \tag{A.7d}$$

Here the kinetic parameters have been scaled such that $\lambda^{-1}$ represents a typical kinetic timescale. We fix the ratio

$$\frac{\mathcal{L}^2}{\mathcal{T}} = \frac{V^2}{\Omega} \tag{A.7e}$$

so that $\mathcal{L}$ and $\mathcal{T}$ are diffusive length and timescales characteristic of the active motion (c.f. [1–3]). We also fix $\mathcal{P}/\mathcal{N} = V/(\mathcal{L}\Omega)$. Substituting (A.7) into (A.5) yields

$$\partial_{\tilde{t}}\tilde{n} = \tilde{D}_t \tilde{\nabla}^2 \tilde{n} - \tilde{v}(\tilde{u})\tilde{\nabla} \cdot \tilde{\mathbf{p}} + \Gamma\left(\tilde{\varepsilon}\partial_{\tilde{u}}^2 \tilde{n} - \partial_{\tilde{u}}\left[\tilde{f}(\tilde{u}, \tilde{c})\tilde{n}\right]\right), \tag{A.8a}$$

$$\partial_{\tilde{t}}\tilde{\mathbf{p}} = \tilde{D}_t \tilde{\nabla}^2 \tilde{\mathbf{p}} - \chi\left(\frac{\tilde{v}(\tilde{u})}{2}\tilde{\nabla}\tilde{n} + \tilde{\omega}_a(\tilde{u})\tilde{\mathbf{p}}\right) + \Gamma\left(\tilde{\varepsilon}\partial_{\tilde{u}}^2 \tilde{\mathbf{p}} - \partial_{\tilde{u}}\left[\tilde{f}(\tilde{u}, \tilde{c})\tilde{\mathbf{p}}\right]\right) + \tilde{\mathbf{M}}[\psi_1], \tag{A.8b}$$

Eqs. (A.8) constitute the dimensionless continuum model of the individual equations of motion (1)–(3).

In order to simplify (A.8), we consider a macroscopic scaling regime in which the timescale of tumbling and reorientation $\Omega^{-1}$ is much faster than both the diffusive timescale $\mathcal{T}$ and kinetic timescale $\lambda^{-1}$. Mathematically, this regime is given by $\chi = \Omega\mathcal{T} \gg 1$ and $\chi \gg \Gamma = \lambda\mathcal{T}$. In this scaling regime, the $\mathcal{O}(\chi)$ terms dominate in (A.8b). Hence $\mathbf{p}$ has the following quasi-steady state to leading order in $\chi$:

$$\mathbf{p} \sim -\frac{v(u)}{2\left[\gamma(u) + D_r(u)\right]}\nabla n, \tag{A.9}$$

where we drop the tilde notation and assume that the higher order moment $\mathbf{M}[\psi_1]$ can be neglected in this regime. The standard reasoning behind dropping the higher order moment (c.f. [1, 2, 4]) is that it is negligible on timescales much longer than the tumbling/reorientation timescale and lengthscales much larger than the persistence length $V/\Omega$. The first condition is satisfied in the regime $\chi \gg 1$. The second is satisfied due to the diffusive scaling (A.7e), i.e. $(V/\Omega)^2 = \mathcal{L}^2/(\Omega\mathcal{T}) \ll \mathcal{L}^2$. Substituting the quasi-steady orientation field (A.9) into the density equation (A.8a) yields

$$\partial_t n = \left(D_t + \frac{v^2(u)}{2\omega(u)}\right)\nabla^2 n + \Gamma\left(\varepsilon\partial_u^2 n - \partial_u\left[f(u, c)n\right]\right). \tag{A.10}$$

Upon reinserting the dimensional quantities (A.7) into (A.10), we obtain the governing equation (4).

In summary, we have derived the simplified governing equation (A.10) by up-scaling the many-body Fokker-Planck equation (A.1) and considering a suitable macroscopic scaling regime. In this scaling regime the timescale of particle tumbling and reorientation is much faster than the kinetic and diffusive timescales. Since we consider the diffusive scaling (A.7e), this is equivalent to considering lengthscales larger than the persistence length, i.e. $\mathcal{L} \gg V/\Omega$. The scaling regime also implies that $\mathcal{P}/\mathcal{N} \ll 1$, which we expect since local order decays on a timescale of $\Omega^{-1}$, as can be seen from the third term on the RHS of (A.5b). We emphasize that no assumption is made about the size of $\Gamma$, so the relationship between the kinetic and diffusive timescales remains general. In our continuum model we consider large populations as are typically present in natural [5] and synthetic [6] biological systems.

Finally, we check that our scaling regime is biologically relevant. We estimate $\chi \approx 900$ and $\Gamma \approx 15$ for run-and-tumble bacteria (no rotational diffusion) with a canonical LuxIR gene-regulatory network and a lengthscale of $\mathcal{L} = 1$mm. These values are in line with the scaling regime $\chi \gg 1$ and $\chi \gg \Gamma$. On a lengthscale comparable to a petri dish ($\mathcal{L} \approx 1$cm), we obtain even more favourable estimates of $\chi \approx 9 \times 10^4$ and $\Gamma \approx 1.5 \times 10^3$. We estimated these values by taking $\lambda \approx 0.01\text{s}^{-1}$ to be the natural decay rate of the LuxR-AI dimer in the prototypical LuxIR system (see $k_-$ in Table S1 of [7]). We also used a tumble rate of $\Omega \approx 0.6\text{s}^{-1}$ and run speed of $V \approx 20\mu$m/s (see the paragraph below Eq. (12) and bottom of page 6 of [8]).

We also note that, as we demonstrate in Section B, our theoretical results can be generalised to the case where the $\mathcal{O}(\Gamma)$ terms in (A.8b) are retained.

### B Asymptotic Analysis of the Linear Stability Problem

In this section we carry out the linear stability analysis in the regime $\varepsilon \ll 1$ in order to derive the dispersion relation in (13) and the instability criterion (14). We additionally perform a linear stability analysis of the system (A.8) and (6)–(9) in the distinguished limit where we retain both the $\mathcal{O}(\chi)$ and $\mathcal{O}(\Gamma)$ terms in (A.8b). The resulting dispersion relation is valid for general relationships between the kinetic and tumbling/reorientation timescales. As such we are able to study how this relationship affects the stability of the system. In the limit where tumbling/reorientation occurs faster than the kinetics, we recover (13) from the main text. The classical MIPS criterion (15) is obtained in the opposite limit where the kinetics are faster than tumbling and reorientation, with the additional assumption of fast AI secretion and decay timescales as clarified below.

We begin by substituting the perturbation (12) into the governing equations (4)-(6) and retaining terms linear in $\eta$ and $C$. After non-dimensionalising with (A.7), we obtain

$$\varepsilon\Gamma\eta'' + \Gamma\left[(u-u_*)\eta\right]' - (\sigma + k^2\mathcal{D})\eta - \Gamma f_c^* n_e' C = 0, \tag{B.11a}$$

$$(\sigma + \beta + k^2 D_c)C - \alpha_0 \int_0^\infty u\eta(u)\,\mathrm{d}u = 0, \tag{B.11b}$$

where $k = |\mathbf{k}|$ and $f_c^* := \partial_c f(u_*, c_*)$. The unperturbed solution is exponentially localised near $u = u_*$ for small $\varepsilon$; we require the same to be true for the perturbation. We therefore look for a WKBJ approximation to $\eta$ in the form

$$\eta(u) = \varepsilon^{-\frac{3}{2}} q(u) \exp\left[-\frac{(u-u_*)^2}{2\varepsilon}\right], \tag{B.12}$$

where the $\varepsilon^{-3/2}$ prefactor is introduced for convenience, so that $q = \mathcal{O}(C)$, as seen in (B.13) below. Substituting (B.12) into (B.11a) gives

$$\varepsilon\Gamma q'' - \Gamma(u-u_*)q' - (\sigma + \mathcal{D}(u)k^2)q = -\frac{\rho_* f_c^* C\Gamma}{\sqrt{2\pi}}(u-u_*), \tag{B.13}$$

where we have used (10) to evaluate $n_e'$. We require $q$ to be polynomially bounded as $u \to 0, \infty$.

We expand the amplitude $q$ in a regular asymptotic power series as

$$q(u) \sim q_0(u) + \varepsilon q_1(u) + \dots. \tag{B.14}$$

The goal now is to calculate $q_0$ and $q_1$ in (B.14) in order to evaluate the integral term in (B.11b). We will find that $q_0(u_*) = 0$, which means that the contributions from $q_0$ and $q_1$ to the integral are comparable, so we cannot ignore $q_1$. Substituting (B.14) into (B.13) and equating coefficients of powers of $\varepsilon$ gives

$$\Gamma(u-u_*)q_0' + \left[\sigma + \mathcal{D}(u)k^2\right]q_0 = \frac{\rho_* f_c^* C\Gamma}{\sqrt{2\pi}}(u-u_*), \tag{B.15a}$$

$$\Gamma(u-u_*)q_1' + \left[\sigma + \mathcal{D}(u)k^2\right]q_1 = \Gamma q_0''. \tag{B.15b}$$

In general, solutions of (B.15) will not be regular at $u_*$. However, since (B.13) is singularly perturbed, any nonsmooth solution of (B.15) will generate a divergent asymptotic expansion (B.14), which in turn will switch on a non-perturbative contribution to $q$ proportional to $\exp((u-u_*)^2/2\varepsilon)$ via Stokes phenomenon [9]. If such a term were present, $q$ would not satisfy the boundary conditions as $u \to 0, \infty$. We therefore deduce that $q$ must be regular at $u_*$.

Since we only require the local behaviour of $q$ in order to apply Laplace's method to the integral in (B.11b), we look for a solution of (B.15) in the form of a regular power series centred at $u = u_*$,

$$q_0(u) = \sum_{\ell=0}^{\infty} c_\ell (u - u_*)^\ell, \tag{B.16}$$

and additionally Taylor expand $\mathcal{D}$ as

$$\mathcal{D}(u) = \sum_{\ell=0}^{\infty} \frac{\mathcal{D}_*^{(\ell)}}{\ell!}(u-u_*)^\ell, \quad \text{with } \mathcal{D}_*^{(\ell)} \equiv \left.\frac{d^\ell}{du^\ell}\right|_{u=u_*} \mathcal{D}(u). \tag{B.17}$$

We then use standard methods to obtain the following recurrence relation for the coefficients:

$$c_\ell = \frac{1}{\sigma + \mathcal{D}_* k^2 + \ell\Gamma}\left[\delta_{\ell 1}\frac{\rho_* f_c^* C\Gamma}{\sqrt{2\pi}} - k^2 \sum_{j=1}^{\ell} \frac{\mathcal{D}_*^{(j)}}{j!} c_{\ell-j}\right], \quad \ell = 1, \ldots, \quad c_0 = 0, \tag{B.18}$$

where $\delta_{ij}$ is the Kronecker delta. We calculate the coefficients in the series for $q_1$ using the same approach. The power series solutions for $q_0$ and $q_1$ read

$$q_0(u) = \frac{\rho_* f_c^* C\Gamma}{\sqrt{2\pi}\,(\sigma + \mathcal{D}_* k^2 + \Gamma)}(u-u_*)\left[1 - \frac{k^2 \mathcal{D}'_*}{\sigma + \mathcal{D}_* k^2 + 2\Gamma}(u-u_*) + \ldots\right], \tag{B.19}$$

$$q_1(u) = -\frac{\sqrt{2}\rho_* f_c^* C\Gamma^2 k^2 \mathcal{D}'_*}{\sqrt{\pi}\,(\sigma + \mathcal{D}_* k^2)(\sigma + \mathcal{D}_* k^2 + \Gamma)(\sigma + \mathcal{D}_* k^2 + 2\Gamma)} + \ldots, \tag{B.20}$$

where we show only the terms required for a leading-order evaluation of (B.11b). Note that $q_0(u_*) = 0$ as anticipated. We can now evaluate the integral in (B.11b) using Laplace's method owing to the fact that $\eta$ is exponentially localised at $u_*$. The leading-order result reads

$$\begin{aligned}\int_0^\infty u\eta(u)du &= \sqrt{\frac{\pi}{2}}\left.[uq_0(u)]''\right|_{u_*} + \sqrt{2\pi}u_* q_1(u_*) + \mathcal{O}(\varepsilon) \\ &= \rho_* f_c^* \Gamma C \frac{\sigma + (\mathcal{D}_* - \mathcal{D}'_* u_*)k^2}{(\sigma + \mathcal{D}_* k^2)(\sigma + \mathcal{D}_* k^2 + \Gamma)} + \mathcal{O}(\varepsilon).\end{aligned} \tag{B.21}$$

Finally, putting the above into (B.11b) and reinserting the dimensional quantities (A.7), yields the dispersion relation for the eigenvalue $\sigma$ given in (13).

In order to derive the instability criterion in (14) from the dispersion relation (13), we put $\sigma = 0$ and rearrange to obtain

$$\frac{\mathcal{D}'_*}{\mathcal{D}_*} = -\frac{1}{u_*}\left[\frac{\left(D_c k^2 + \beta\right)\left(\mathcal{D}_* k^2 + \lambda\right)}{\alpha_0 \rho_* f_c^*} - 1\right], \tag{B.22}$$

where it can be shown that the right side is strictly negative. Equation (B.22) is the criterion for instability for a given wavenumber $k$. The right side of (B.22) is a decreasing function of $k^2$ so as $\mathcal{D}'_*$ is decreased from zero, the first mode to become unstable is the one corresponding to the smallest allowed value of $k$. Thus we obtain (14) by using $k = \pi/L_m$, where $L_m$ is the largest side length of the domain.

In the remainder of this section we generalise the above linear stability analysis to the case where we drop the assumption that the tumbling and reorientation timescale is much faster than the kinetic timescale. Specifically, we drop the assumption $\chi \gg \Gamma$ and therefore retain the $\mathcal{O}(\Gamma)$ terms in (A.8), rather than using (A.9). Our governing equations for the linear stability analysis consist of (A.8) and (6)–(9). The analysis utilizes similar WKBJ techniques as shown above for the simplified system. We therefore provide only an outline of the analysis and flag the important differences. Finally, we briefly show how the classic MIPS criterion (15) arises from the stability analysis in the regime $\Gamma \gg \chi$ and fast AI secretion and decay timescales (clarified below).

Our starting point is (A.8). We keep the assumption that $\chi \gg 1$, so that $\mathcal{O}(1)$ terms can be neglected. As such, our analysis is valid for $\Gamma$ ranging from $\mathcal{O}(1)$ to much larger than $\chi$. The time derivative, translational diffusion, and higher-order moment terms in (A.8b) are all $\mathcal{O}(1)$ and are therefore negligible. Retaining both the $\mathcal{O}(\chi)$ and $\mathcal{O}(\Gamma)$ terms in (A.8b) and dropping the tilde notation we obtain

$$\partial_t n = -v(u)\nabla\cdot\mathbf{p} + \Gamma\left(\varepsilon\partial_u^2 n - \partial_u\left[f(u,c)n\right]\right), \tag{B.23a}$$

$$0 = -\chi\left(\frac{v(u)}{2}\nabla n + \omega_a(u)\mathbf{p}\right) + \Gamma\left(\varepsilon\partial_u^2\mathbf{p} - \partial_u\left[f(u,c)\mathbf{p}\right]\right). \tag{B.23b}$$

The simplified system (A.10) is recovered from (B.23) at leading order in $\chi$ when $\chi \gg \Gamma$, physically corresponding to fast tumbling/reorientation timescales compared with kinetic timescales.

The homogeneous steady-state of the governing equations (B.23) is now given by $\mathbf{p} = 0$, $n = n_e(u)$, and $c = c_*$, where $n_e$ and $c_*$ are defined in (10) (in dimensional quantities). As before we introduce small perturbations of the form

$$n(\mathbf{x},u,t) = n_e(u) + \eta(u)e^{i\mathbf{k}\cdot\mathbf{x}+\sigma t}, \quad \mathbf{p}(\mathbf{x},u,t) = \mathbf{P}(u)e^{i\mathbf{k}\cdot\mathbf{x}+\sigma t}, \quad c(\mathbf{x},t) = c_* + Ce^{i\mathbf{k}\cdot\mathbf{x}+\sigma t}. \tag{B.24}$$

The perturbations are required to be exponentially localised near $u = u_*$. As such, we look for a WKBJ solution to $\eta$ and $\mathbf{P}$ of the form

$$\eta(u) = \varepsilon^{-3/2}q(u)\exp\left[-\frac{(u-u_*)^2}{2\varepsilon}\right], \quad \mathbf{P}(u) = \varepsilon^{-3/2}i\mathbf{k}s(u)\exp\left[-\frac{(u-u_*)^2}{2\varepsilon}\right]. \tag{B.25}$$

Substituting (B.24)–(B.25) into (B.23) and (6) gives the following amplitude equations for $q$ and $s$

$$\varepsilon\Gamma q'' - \Gamma(u-u_*)q' - \sigma q + vk^2 s = -\frac{\rho_* f_c^* C\Gamma}{\sqrt{2\pi}}(u-u_*), \tag{B.26a}$$

$$\varepsilon\Gamma s'' - \Gamma(u-u_*)s' - \chi\omega_a s - \frac{\chi v q}{2} = 0. \tag{B.26b}$$

We require $q$ and $s$ to be polynomially bounded as $u \to 0, \infty$ as boundary conditions. In the limit $\varepsilon \ll 1$, we expand the amplitudes $q$ and $s$ in a regular asymptotic power series (c.f. Eq. (B.14)). The terms in this expansion are then calculated via standard power series methods. This lengthy but straightforward calculation results in the dispersion relation

$$\sigma + D_c k^2 + \beta - \alpha_0 f_c^* \rho_* g(\sigma) = 0, \tag{B.27a}$$

where $g$ is given by

$$g(\sigma) = \frac{\chi\Gamma\left[\sigma + (\mathcal{D}_* - u_*\mathcal{D}'_*)k^2\right] + \frac{\Gamma^2}{\omega_{a*}}\left[\sigma + \mathcal{D}_* k^2\left(1 - u_*\frac{v'_*}{v_*}\right)\right]}{(\sigma + \mathcal{D}_* k^2)\left[(\sigma+\Gamma)\left(\chi + \frac{\Gamma}{\omega_{a*}}\right) + \chi\mathcal{D}_* k^2\right]}, \tag{B.27b}$$

where subscript '$*$' denotes evaluation at $u_*$. Eq. (B.27) constitutes a third-order polynomial equation from which we can in principle determine $\sigma$. Any solution $\sigma$ that has $\mathrm{Re}(\sigma) > 0$ leads to an instability in the uniform steady-state.

We demonstrate that our stability result for the general system (A.8) is consistent with the result for the simplified system (A.10) considered in the main text. Specifically, we show that (B.27) reduces to (13) in the expected limit where tumbling and reorientation occur much faster than the kinetics, i.e. $\chi \gg \Gamma$. In order to show this, we simply retain the dominant $\mathcal{O}(\chi)$ terms in both the numerator and denominator of (B.27b). Then to leading–order in $\chi$, Eq. (B.27a) becomes

$$\sigma + D_c k^2 + \beta - \alpha_0 f_c^* \rho_* \Gamma \frac{\sigma + (\mathcal{D}_* - u_* \mathcal{D}'_*) k^2}{(\sigma + \mathcal{D}_* k^2)\,(\sigma + \mathcal{D}_* k^2 + +\Gamma)} = \mathcal{O}(\chi^{-1}). \tag{B.28}$$

Upon reinserting the dimensional quantities (A.7), we recover (13).

We now briefly demonstrate how the classic MIPS criterion (15) arises from the dispersion relation (B.27). This requires considering the limit where the kinetics occur much faster than tumbling and reorientation ($\Gamma \gg \chi$) and where AI decay and secretion occur faster than AI diffusion ($\beta \sim \alpha_0 \gg D_c = \mathcal{O}(1)$). We recall that our derivation of (B.27) already assumes fast tumbling and reorientation compared to cell diffusion ($\chi \gg 1$). Using the above assumptions, we retain the $\mathcal{O}(\Gamma^2)$ terms in both the numerator and denominator of (B.27b) (note that $g(\sigma) = \mathcal{O}(1)$) and neglect the first two terms on the LHS of (B.27a). This leads to

$$\sigma \sim -\frac{\mathcal{D}_* k^2 \left[\beta - \alpha_0 f_c^* \rho_* \left(1 - u_* \frac{v'_*}{v_*}\right)\right]}{\beta - \alpha_0 f_c^* \rho_*}. \tag{B.29}$$

Using the definition of $f$ in (5), it can be shown that the denominator is strictly positive. Instability thus requires

$$\beta - \alpha_0 f_c^* \rho_* \left(1 - u_* \frac{v'_*}{v_*}\right) < 0. \tag{B.30}$$

Using the chain rule, we exchange the derivative $v'(u_*)$ for a derivative with respect to $\rho$. This leads to

$$\left.\frac{dv}{d\rho}\right|_{\rho_*} < -\frac{1}{\rho_*}, \tag{B.31}$$

which is the well-known MIPS criterion [10]. We note that (B.27) is effectively a cubic equation for $\sigma$ that is singular in the asymptotic limit $\Gamma \gg \chi$. Accordingly, we obtain only one root. It can be shown using standard methods that the remaining two roots have strictly negative real part and thus do not contribute an instability.

### C Calculation of the Hopf Bifurcation Point

In this section we derive the instability criterion (16) from the dispersion relation (13). At a Hopf bifurcation the eigenvalue $\sigma$ is purely imaginary. Substituting $\sigma = i\nu$, with $\nu \in \mathbb{R}$, in (13) the real and imaginary components are found to satisfy

$$\nu^2 - [\omega_0 \omega_\lambda + \omega_\beta(\omega_0 + \omega_\lambda) - \alpha f_c^* \rho_*] = 0, \tag{C.32a}$$

$$(\omega_0 + \omega_\lambda + \omega_\beta)\nu^2 + \omega \omega_\lambda \omega_\beta - \alpha f_c^* \rho_* (\omega_0 - \mathcal{D}'_* u_* k^2) = 0. \tag{C.32b}$$

The two equations for $\nu$ are consistent if, and only if,

$$\omega_0 \omega_\lambda + \omega_\beta(\omega_0 + \omega_\lambda) - \alpha f_c^* \rho_* = \frac{\omega_0 \omega_\lambda \omega_\beta - \alpha f_c^* \rho_* (\omega_0 - \mathcal{D}'_* u_* k^2)}{\omega_0 + \omega_\lambda + \omega_\beta}. \tag{C.33}$$

After rearranging for $\mathcal{D}'_*$, and a little algebra, we obtain the instability criterion (16).

### D Reduction to Classical MIPS

The aim in this section is to clarify the conditions under which our system is formally equivalent to a system of physically interacting particles. We have shown in Section B that we recover the well-known instability criterion (B.31) in the regime of

1. fast AI secretion and decay timescales as compared to AI diffusion timescales ($\beta \sim \alpha_0 \gg D_c$)
2. fast reaction kinetics with respect to the tumbling and reorientation timescales ($\Gamma \gg \chi$)
3. fast tumbling and reorientation with respect to cell diffusion ($\chi \gg 1$)

In the same timescale regime, we show that the time–dependent system (A.8) is dynamically equivalent to a system of physically interacting particles. At the continuum level, physical inter–particle interactions generally appear as a density dependence of the motility parameters [1, 11, 12]. For pure ABPs, the self–propulsion speed is approximately linear. We describe a special case where the gene-regulated motility in our model reduces to a linear density-dependent motility, thereby effectively representing repulsive ABPs.

We begin by considering (A.8) in the regime characterised by 1.–3. above. We also set $\varepsilon = D_t = 0$ for simplicity. We retain a general $D_r(u)$ and $\gamma(u)$ in order to demonstrate that non-constant reorientation and tumble rates cannot trigger MIPS in this regime. Eq. (A.8) thus becomes

$$\partial_t n = -v(u)\nabla \cdot \mathbf{p} - \Gamma \partial_u \left[f(u,c)n\right], \tag{D.34a}$$

$$\partial_t \mathbf{p} = -\chi \left[\frac{v(u)}{2}\nabla n + \omega_a(u)\mathbf{p}\right] - \Gamma \partial_u \left[f(u,c)\mathbf{p}\right] + \mathbf{M}[\psi_1], \tag{D.34b}$$

where we again drop the tilde notation.

In the regime $\Gamma \gg \chi \gg 1$ the reaction term dominates in both (D.34a) and (D.34b), leading to a highly localised distribution of internal chemical:

$$n(\mathbf{x},u,t) \sim \rho(\mathbf{x},t)\delta(u - u_*(\mathbf{x},t)), \tag{D.35a}$$

$$\mathbf{p}(\mathbf{x},u,t) \sim \mathbf{Q}(\mathbf{x},t)\delta(u - u_*(\mathbf{x},t)), \tag{D.35b}$$

$$f(u_*,c) = 0. \tag{D.35c}$$

Assumption 1. above implies that Eq. (6) for $c$ reduces to

$$\beta c(\mathbf{x},t) \sim \alpha(u_*(\mathbf{x},t))\rho(\mathbf{x},t). \tag{D.35d}$$

Eqs. (D.35c) and (D.35d) constitute an algebraic system that, in principle, can be solved for $u_*$ and $c$ in terms of $\rho$. We therefore make use of the shorthand $u_*(\rho) := u_*(\rho(\mathbf{x},t))$. Next, we integrate (D.34b) over $u$ and make use of Assumption 2. above, which yields

$$\omega_a(\rho)\mathbf{Q}(\mathbf{x},t) = -\frac{1}{2}\nabla\left(\rho\int_0^\infty v(u)\delta(u - u_*(\mathbf{x},t))\,\mathrm{d}u\right) = -\frac{1}{2}\nabla(v(\rho)\rho), \tag{D.36}$$

where $v(\rho) = v(u_*(\rho))$ and $\omega_a(\rho) = \omega_a(u_*(\rho))$ and we have assumed that the higher-order angular moment term can be neglected. We have also used the fact that

$$\int_0^\infty \omega_a(u)\mathbf{p}\,\mathrm{d}u = \mathbf{Q}(\mathbf{x},t)\int_0^\infty \omega_a(u)\delta(u - u_*(\mathbf{x},t))\,\mathrm{d}u = \omega_a(\rho)\mathbf{Q}(\mathbf{x},t). \tag{D.37}$$

Integrating (D.34a) over $u$ and substituting (D.36) leads to

$$\partial_t \rho = \nabla \cdot \left[\left(\frac{v^2}{2\omega_a} + \frac{vv'\rho}{2\omega_a}\right)\nabla\rho\right], \tag{D.38}$$

where $v' = v'(\rho)$. Eq. (D.38) has a linear instability when $v'/v < -\rho^{-1}$, consistent with (15) and (B.31). Note also that variations in $\omega_a$ alone cannot trigger MIPS, consistent with previous literature [4, 10]

In the very-fast-chemical-timescale limit ($\Gamma \gg \chi \gg 1$), the density dependence of the motility $v(u_*(\rho_*))$ will in general be non-linear in the density. A linear motility arises as a special case when

1. there is no positive feedback in the QS system, corresponding to a constant secretion rate $\alpha(u) = \alpha_*$,
2. the kinetics are linear in both the internal concentration $u$ and the AI concentration $c$ (c.f. [13]), e.g. $f(u, c) = a + Lc - \lambda u$
3. $v(u)$ is linear in the internal chemical $u$.

Assumptions 1 and 2 imply that (D.35c) and (D.35d) reduce to

$$\beta c = \alpha_* \rho(\mathbf{x}, t), \quad a + Lc - \lambda u_* = 0. \tag{D.39}$$

Then assumption 3 gives a linear $v(\rho)$

$$v(u_*(\rho)) = v_0 - \zeta u_*(\rho) = \bar{v}_0 - \bar{\zeta}\rho(\mathbf{x}, t), \tag{D.40}$$

where we absorb constants of proporitionality into the coefficients.

### E Numerical Solution of the Governing Equations using oomph-lib

In this section we describe the general implementation of the numerical solution of the governing equations (4)–(9) within the open-source finite-element library oomph-lib [14]. Broadly speaking, we use a Galerkin finite element method with quadratic Lagrange elements with a BDF2 time stepping scheme. However, there are two main features of the governing equations that prevent a 'standard' implementation. The first is that the AI concentration $c$ depends on the structured cell density non-locally through the integral term in (6). This term is non-local in the $u$ coordinate only. Secondly, equations (4) and (6) are defined on domains of different dimension due to the $u$ dependence. The implicit BDF2 scheme uses a finite-difference approximation for the Jacobian. For the steady problem, we use an analytical Jacobian in order to compute steady-state solutions near a bifurcation point. The steady problem is more complicated and we discuss it further at the end of this section. The parameter values used in the numerical simulations can be found in Table S1.

We accommodate the non-local term in (6) by representing it with an auxiliary field $\bar{u}(\mathbf{x})$, defined as

$$\bar{u}(\mathbf{x}) = \int_0^\infty u n(u, \mathbf{x}, t)\, \mathrm{d}u. \tag{E.41}$$

This transforms the governing equations (4) and (6) into two local equations coupled to one non-local algebraic equation (c.f [15]). Recall that we use $\alpha(u) = \alpha_0 u$ in (6). Eq. (E.41) for $\bar{u}$ is already in weak form in which $\mathbf{x}$ is a parameter.

In order to deal with the difference in domain dimension of (4) and (6), we extend the domain of (6) into the $u$ direction. As such, we use test functions defined on $\Omega \times \mathbb{R}_+$ when deriving a weak form of the governing equations. We use a multi-domain approach in which the governing equations for $c$ and $n$ (in weak form) are solved on copies of the same domain with different discretisations. In other words, $n$ and $c$ are restricted to different finite dimensional subspaces in the numerical solution. In both cases we use piece-wise quadratic basis functions defined on rectangular elements. We note that this choice is ideal for evaluating the integral in (E.41) since it leads to a straightforward application of Simpson's rule, which is exact for our choice of basis functions. Clearly we must look for solutions $c(\mathbf{x}, t)$ that are independent of $u$, so we simply use a mesh that is one element wide in the $u$ direction[a)] and verify *a posteriori* that the numerical solution for $c$ is $u$-independent. The mesh for $n$ is much finer in the $u$ direction, but otherwise contains the same number of elements in the $\mathbf{x}$ direction as the mesh for $c$. A separate rectangular mesh is used for $\bar{u}$ which, for simplicity, occupies only $\Omega$.

In addition to the above complications, the steady-state problem has the added difficulty that the uniform cell density $\rho_*$ is effectively a free parameter. In the time-dependent case, it is determined from the initial condition owing to the fact that the total number of cells in the population is conserved, i.e.

$$\rho_* = \frac{1}{|\Omega|} \int_\Omega \int_0^\infty n(u, \mathbf{x}, t)\, \mathrm{d}u \mathrm{d}\mathbf{x}, \tag{E.42}$$

where $|\Omega|$ is the volume of the domain $\Omega$. Eq. (E.42) is obtained by integrating (4) over $u$ and $\mathbf{x}$. The parameter $\rho_*$ must be specified in the numerical solution somehow. Our approach here is to use the concept of a 'hijacked' element in which we replace the elemental residual of one element in the mesh with a different residual that enforces the population constraint [14]. This effectively removes the extra degree of freedom associated with the free parameter $\rho_*$. The equation for the traded residual on the hijacked element is given by rearranging (E.42) such that the right side is equal to zero. The implementation involves defining a new mesh of elements that calculate the elemental contributions to the integral in (E.42), as well as one additional element that assembles the contributions and evaluates the traded residual. A similar application of the hijacking method can be found in the online documentation for oomph-lib [16].

We use the numerical solution of the steady-state problem to calculate the blue and red lines in the bifurcation diagram shown in figure 2. The cell density $\rho(x)$ is calculated by integrating the steady-state numerical solution $n(u, x)$ over $u$ using an exact quadrature for the finite element basis. We then calculate the difference between the global maximum and global minimum of the cell density and repeat the process using pseudo arc–length continuation in the parameter $\mathcal{D}'_*$.

All numerical calculations use a specified piece-wise linear effective diffusion coefficient. In the case where the internal chemical suppresses motility (case I in the main text), we use

$$\mathcal{D}(u) = \begin{cases} \mathcal{D}'_*(u - u_*) + \mathcal{D}_*, & 0 \le u < u_c, \\ \mathcal{D}_\infty, & u \ge u_c. \end{cases} \tag{E.43}$$

When the internal chemical promotes motility (case II), we use

$$\mathcal{D}(u) = \begin{cases} \mathcal{D}_0, & 0 \le u < u_c, \\ \mathcal{D}'_*(u - u_*) + \mathcal{D}_*, & u > u_c. \end{cases} \tag{E.44}$$

Here $u_*$ is the steady-state mean internal concentration defined in (11) while $\mathcal{D}_*$ and $\mathcal{D}'_*$ are parameters. In both cases, $u_c$ is selected such that $\mathcal{D}$ is continuous.

### F Internal Mean-Field Model

We develop a mean-field model with respect to the internal chemical $u$ in order to quantify the difference between an internal mean-field approach and a gene-structured approach, highlighting

[a)] We use two elements wide in 2 spatial dimensions.

| Symbol | Description | Value (Case I) | Value (Case II) |
|---|---|---|---|
| $\mathcal{D}_*$ | Effective motility coefficient in the uniform state | 0.5 | 0.5 |
| $\mathcal{D}'_*$ | Motility response in the uniform state | Variable $(-5, -0.19)$, see figures 1–2 | Variable $(3, 16)$, see figure 3 |
| $\mathcal{D}_0$ | Motility coefficient at zero internal concentration | 0.1 | – |
| $\mathcal{D}_\infty$ | Motility coefficient at large internal concentrations | – | 0.1 |
| $\varepsilon$ | Strength of stochasticity in the GRN kinetics | 0.04 | 0.001 |
| $a$ | Base production rate of TF in GRN kinetics | 0.3 | 0.3 |
| $L$ | Saturation production rate of positive feedback in GRN kinetics | 10 | 15 |
| $K$ | Threshold activation concentration of positive feedback in GRN kinetics | 10 | 15 |
| $\lambda$ | Decay rate of TF | 2 | 1 |
| $D_c$ | Diffusion coefficient of AI | 0.4 | 0.4 |
| $\beta$ | Decay rate of AI | 4 | 5 |
| $\alpha_0$ | Secretion rate of AI | 12.5 | 6.25 |
| $\rho_*$ | Uniform cell density | 1 | 1 |
| $N_x$ | Number of elements in the $x$ coordinate for numerical calculations | 25 (2D), 50 (1D) | 50 |
| $N_y$ | Number of elements in the $y$ coordinate for numerical calculations | 25 | – |
| $N_u$ | Number of elements in the $u$ coordinate for numerical calculations | 100 (2D), 300 (1D) | 200 |

Table S1: List of parameters and their values used in the numerical calculations for figures 1–3 in the main text. All quantities are dimensionless.

the advantages of the latter. An internal mean-field model implicitly assumes that cells have locally identical internal concentrations. That is, the mean internal concentration contains the full information of the distribution of internal concentration if, and only if, all internal concentration are equal to the mean. Mathematically, this corresponds to a highly localised (in $u$) structured density $n(u, \mathbf{x}, t)$.

We derive an internal mean-field model which satisfies the zeroth and first order moments of the governing equation (4) by assuming that $n$ is highly localised in $u$, i.e.

$$n(u, \mathbf{x}, t) = \delta(u - u_*(\mathbf{x}, t))\rho(\mathbf{x}, t). \tag{F.45}$$

Here $\delta(\cdot)$ is the Dirac delta function and $u_*(\mathbf{x}, t)$ is the mean internal concentration to be determined. The zeroth moment of $n$ is equal to $\rho$ and the first moment $\bar{u}(\mathbf{x}, t)$ is related to $\rho$ and $u_*$ through

$$\bar{u}(\mathbf{x}, t) := \int_0^\infty un(u, \mathbf{x}, t)du = \rho(\mathbf{x}, t)u_*(\mathbf{x}, t). \tag{F.46}$$

In order to obtain governing equations for $\rho$ we calculate the zeroth moment of (4) by integrating over $u$ to obtain

$$\partial_t \rho = \int_0^\infty \mathcal{D}(u)\nabla^2 \left[\delta(u - u_*(\mathbf{x}, t))\rho(\mathbf{x}, t)\right] du, \tag{F.47}$$

where we have set $\varepsilon = 0$ for simplicity. Since $\mathcal{D}$ is independent of $\mathbf{x}$, it can be brought inside the Laplacian. We then use the sifting property of the delta function to obtain

$$\partial_t \rho = \nabla^2 \left[\mathcal{D}(u_*)\rho\right]. \tag{F.48a}$$

The governing equation for $u_*$ is found in a similar fashion by considering the first moment of (4). The result is

$$\partial_t(\rho u_*) = \nabla^2 \left[\mathcal{D}(u_*)\rho u_*\right] + f(u_*, c)\rho, \tag{F.48b}$$

where $f$ is given by (5) and we have used (F.46) to replace $\bar{u}$ in favour of $u_*$. Finally, we simply rewrite (4) in terms of the mean-field quantities as

$$\partial_t c = \nabla^2 c - \beta c + \alpha_0 u_* \rho. \tag{F.48c}$$

Eqs. (F.48), together with no-flux boundary conditions, constitute our internal mean-field model.

The internal mean-field equations agree with the structured model near the homogeneous steady-state, as is to be expected since the uniform steady-state of (F.48a)–(F.48c) is consistent with (10)–(11). Additionally, the two instability criteria, obtainable via linear stability analysis of (F.48a)–(F.48c), are consistent with (14) and (16). However the spatially heterogeneous MIPS patterning is quantitatively different in the two approaches. These differences are illustrated in figure S1 below.

We expect that an internal mean-field approach will achieve best agreement with a structured model for the simplistic kinetics considered herein. If the kinetics are non-linear in $u$, the moment equations (F.48a) and (F.48b) will depend on higher order moments so that moment closure techniques are required to obtain a closed system. We expect this internal mean-field approach to be much less accurate with more realistic kinetics (e.g. kinetics with multiple equilibria).

To produce Figure S1, the structured equations (4)–(6) and the internal mean-field equations (F.48a)–(F.48c) are solved numerically with oomph-lib [14]. We use a Galerkin finite element method with quadratic Lagrange elements and BDF2 time-stepping. In order to compare the results with the gene-structured model, we use the same parameter set (see the third column of Table S1), with the exception that $\varepsilon = 0$ and we use a slightly modified diffusion coefficient $\mathcal{D}$ and higher spatial resolution ($N_x = 1000$). The reason that $\mathcal{D}$ is adjusted is that the coefficients of the highest order derivatives in (F.48a)–(F.48b) involve terms containing $\mathcal{D}'$, which is discontinuous if (E.43) or (E.44) is used. As such, we introduce a small cubic regularisation near $u_c$ in (E.43) so that $\mathcal{D}'$ is continuous.

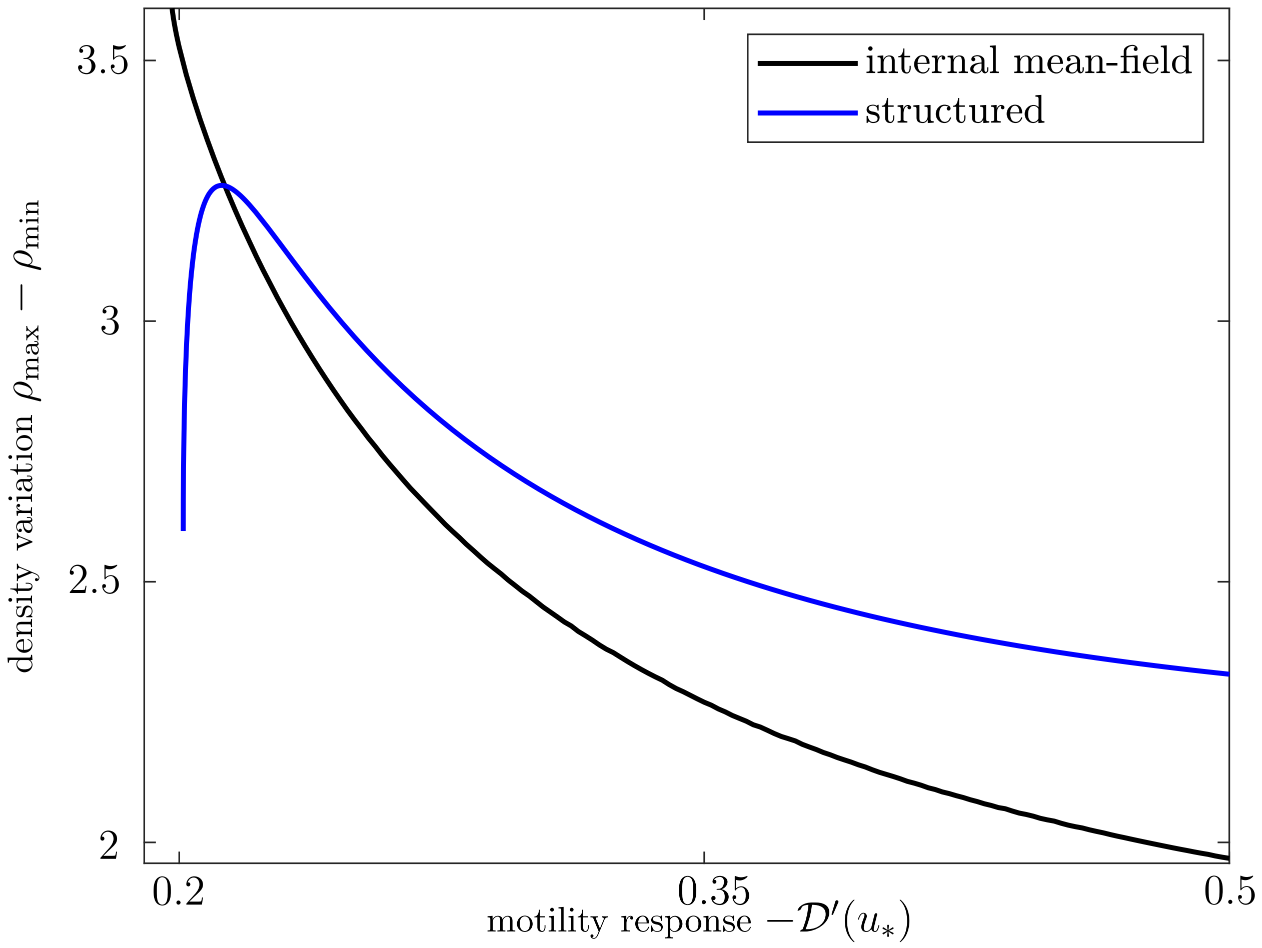


Figure S1: Stable MIPS patterned branches of the 1D bifurcation diagrams for the structured model in the main text and internal mean-field model (F.48).
